# Bayesian Estimation of Muscle Mechanisms and Therapeutic Targets Using Variational Autoencoders

**DOI:** 10.1101/2024.05.08.593035

**Authors:** Travis Tune, Kristina B Kooiker, Jennifer Davis, Thomas Daniel, Farid Moussavi-Harami

## Abstract

Cardiomyopathies, often caused by mutations in genes encoding muscle proteins, are traditionally treated by phenotyping hearts and addressing symptoms post irreversible damage. With advancements in genotyping, early diagnosis is now possible, potentially introducing earlier treatment. However, the intricate structure of muscle and its myriad proteins make treatment predictions challenging. Here we approach the problem of estimating therapeutic targets for a mutation in mouse muscle using a spatially explicit half sarcomere muscle model. We selected 9 rate parameters in our model linked to both small molecules and cardiomyopathy-causing mutations. We then randomly varied these rate parameters and simulated an isometric twitch for each combination to generate a large training dataset. We used this dataset to train a Conditional Variational Autoencoder (CVAE), a technique used in Bayesian parameter estimation. Given simulated or experimental isometric twitches, this machine learning model is able to then predict the set of rate parameters which are most likely to yield that result. We then predict the set of rate parameters associated with twitches from control mice with the cardiac Troponin C (cTnC) I61Q variant and control twitches treated with the myosin activator Danicamtiv, as well as model parameters that recover the abnormal I61Q cTnC twitches.

**SIGNIFICANCE:** Machine learning techniques have potential to accelerate discoveries in biologically complex systems. However, they require large data sets and can be challenging in high dimensional systems such as cardiac muscle. In this study, we combined experimental measures of cardiac muscle twitch forces with mechanistic simulations and a newly developed mixture of Bayesian inference with neural networks (in autoencoders) to solve the inverse problem of determining the underlying kinetics for observed force generation by cardiac muscle. The autoencoders are trained on millions of simulations spanning parameter spaces that correspond to the mechanochemistry of cardiac sarcomeres. We apply the trained model to experimental data in order to infer parameters that can explain a diseased twitch and ways to recover it.

## INTRODUCTION

In muscle, force is generated by the interaction of two overlapping filaments, the myosin containing thick filaments and the actin containing thin filaments (Fig. 1). The contractile process is regulated by troponin-tropomyosin complexes spaced along each thin filament. In the presence of calcium ions, these complexes undergo a conformational change, allowing myosin motors to attach to actin binding sites on the thin filament. Upon binding, myosin motors generate force (1). This complex series of chemo-mechanical events can be modeled as state transitions in the regulatory proteins in the thin filament and energy (ATP) transformations of the myosin crossbridges extending from the thick filament (Fig. 1) (2–6). Both genetic and acquired muscle disorders can alter or interrupt the chemo-mechanical state transitions of contractions, with several of the above studies using computational simulations to connect estimates of state transition rates to observed mechanical behaviors of abnormal muscle contraction. These approaches generally fall into the category of forward models – that is they generate predictions of muscle contractile behavior based on the best possible estimates of the kinetics of state transitions as well as the mechanics and geometry of the contractile lattice.

**Figure 1:**
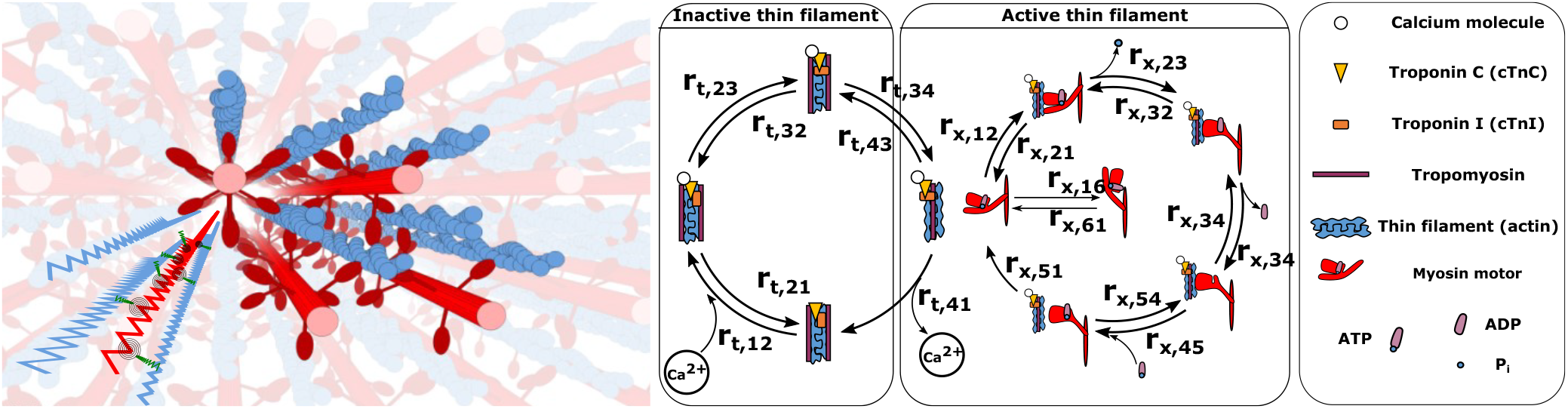
Left: 3D rendering of a sarcomere’s lattice showing the thick (red) and thin filaments (blue) as well as a superimposed spring network to give a sense of the model’s geometry. Following the geometry specified in (7) we model the filaments as a network of springs for the thick and thin filaments, with crossbridges, each consisting of a torsional and linear spring. Right: The rate transition diagram for the thick and thin filaments, indicating the four thin filament rates (labeled *r*_*t*_) and the six myosin motor rates (labeled *r*_*x*_), as well as a table indicating the key biological elements. Crossbridges are modeled as a torsional and linear spring.

Recently, advances in machine learning and optimization methods allow us to address the connection between the underlying mechanochemistry of muscle and the emergent behavior as an inverse problem. That is, given the observed contractile behavior, what underlying kinetics, mechanics, and geometry best explain the observation? Answering this question through experimental efforts alone is both time consuming and challenging. Simulations, on the other hand, can allow us to both explore a vast parameter space as well as estimate the effect of different underlying parameters. However, even then, there are significant challenges regarding selection of rate constants or other model parameters at baseline or with particular diseased states. For one thing, because we are exploring a high dimensional parameter space, our system is under-determined, and there are likely many different combinations of rate constants which could correspond to indistinguishable outcomes. Also, because many current simulations are based on Monte-Carlo methods (3, 6), random variation in both the simulation and experimental data can make predictions difficult. Solving the inverse problem requires a very large data set that spans many combinations of rate constants. A common way to approach inverse problems is by reformulating them as a probabilistic inference problem using Bayes’ Theorem. Rather than seeking a particular solution to a problem, we treat the solution as a random variable, and attempt to quantify our uncertainty in its value. This allows for incorporating uncertainty, whether from noise or the under-determined nature of the model in question, into our predictions. A challenge of Bayesian inference in high dimensional data is slow convergence, which can be improved using machine learning techniques.

Previously, a machine learning architecture called Conditional Variational Autoencoders (CVAE) have been used in other contexts in order to solve high dimensional noisy inverse inference problems (8). Auto encoders work by learning a distribution over an abstract latent space of an input, and then learning to reconstruct the input from the reduced representation. In order to solve the inverse problem of rate parameters that describe a particular twitch, we first generated a very large set of twitches using our spatially explicit half sarcomere muscle model (2, 7, 9). In this model, the thick and thin filaments are comprised of a series of springs with stiffnesses determined experimentally. Our model allows us to specify the geometric configuration of filaments, myosin motors, and actin binding sites, and allows us to alter the rate kinetics of motors or binding sites (6, 7, 10), or even a subset of the population according to any spatial distribution. We first trained the model to reduce and reconstruct the distribution describing the vector of rate parameters 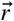 using both the vector (in time) of a twitch, *f*, and 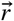. We then tested the model by giving it only information on *f* alone. The posterior distribution 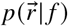 represents our belief that a set of rates 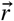 will result in an observation *f*, conditioned on our specific dataset, generated using our spatially explicit model. We are able to successfully train that model to predict the probability of a given rate constant resulting in a particular twitch. We then apply the model to a set of experimental mouse cardiac twitches and predict possible combinations of rate constants that can produce a mouse genetic dilated cardiomyopathy (DCM) twitch or mouse twitches in muscle treated with a myosin activator. Lastly, we predict a combination of parameters that can best recover the abnormal DCM twitch. We show feasibility and validation of using machine learning techniques to infer underlying mechanism of diseased muscle and best ways to correct them.

## MATERIALS AND METHODS

### Animals

All experiments followed protocols approved by the University of Washington Institutional Animal Care and Use Committee according to the “Guide for the Care and Use of Laboratory Animals” (National Research Council, 2011). We used a previously published transgenic mouse model of genetic cardiomyopathy that over expresses the calcium desensitizing I61Q cTnC (cardiac troponin C) variant. The transgenic mice have about 50% cTnC replaced with the I61Q cTnC variant and have cardiac chamber dilation and reduced contractility compared to wild type mice (11, 12). Control and I61Q cTnC male and female mice between the ages of 4-6 months were used to collect experimental cardiac twitch data. Danicamtiv treated twitches are imported from our previously published study (12).

### Experimental mouse cardiac twitch force measurements

Control and I61Q cTnC data presented here were newly collected for this project. Intact papillary muscles were dissected from the right ventricle of mouse hearts, secured between two aluminum t-clips (Aurora Scientific, Ontario, Canada) and mounted into the IonOptix Intact Muscle Chamber (IonOptix, Westwood, MA) between a force transducer and a length controller. The papillary was then submerged in an experimental chamber and was continuously perfused with modified Krebs buffer (1.8 *mM* CaCl) at 33 °C aerated with a 95/5 percent oxygen/carbon dioxide gas mixture to maintain physiological pH. Twitches were generated after field stimulation at 1 Hz with oscillating polarity. The initial length was set to just above slack. After pacing for about 20 minutes at 1 Hz, papillary muscles were stretched to an optimal length where peak twitch no longer increases for data acquisition. Width and thickness were measured for each muscle preparation to determine cross-sectional area, and all twitches are reported as stress 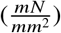. Data was analyzed using the IonOptix IonWizard software. Twitches were averaged over a 30 second recording with ten individual twitches for each preparation exported. The Danicamtiv data - and its corresponding control data - shown here were collected on a different set up and previously published in (12).

### Spatially explicit half-sarcomere model

We began with the spatially explicit model as described in our previous publications (2, 4, 6, 7, 9, 10, 13, 14). In this model, we define a 3D lattice of thick and thin filaments, each composed of a series of elastic elements in which the nodes represent myosin crossbridges and actin binding sites respectively. We have relied on these prior publications above to set values for the parameters that follow below. For example, the stiffness of *k*_*thick*_ (2020 *pn / nm*) and *k*_*thin*_ (1743 *pn / nm*) are determined by measuring the stiffness of entire thick and thin filaments by a combination of microscopy and x-ray diffraction and using the repeat distances of 38.7 and 43 *nm* to scale the stiffness of each segment of the two filament types (2, 15, 16). Titin is included, and is modeled as a passive exponential spring as described in (7, 10):

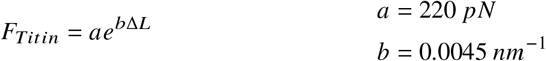

Crossbridges themselves are composed of a torsional and linear spring, situated at the nodes of the thick filament. The three state model described in previous articles consisted of a single free state, a loosely bound state (pre-powerstroke), and a tightly bound state (post-powerstroke). The stiffness of the crossbridge states and the zero points for each spring for each state are determined by electron tomography and x-ray diffraction (6, 17, 18). Force generated by the crossbridges is the spring force of the torsional and linear spring system composing the crossbridges. Here, *r* and *θ* represent the polar distance from a myosin head to its nearest binding site, and *k*_*r*_ and *k* _*θ*_ refer to the stiffness of the linear spring element (the globular domain) and torsional stiffness (the converter domain), and the subscripts (W,S) refer to the weak and strong states, with the change in the spring equilibrium point being what generates force. Energies are given in units of KT.

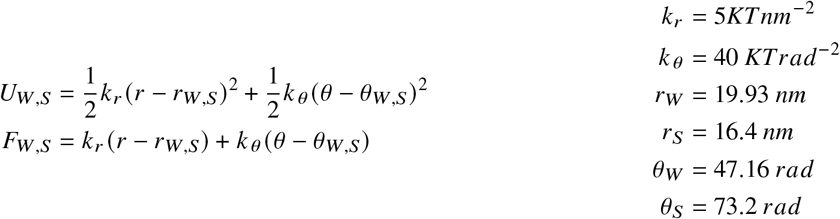

In order to more easily connect to experimental measurements of state transitions and other models, we have expanded our previous spatially explicit model to incorporate additional states in the crossbridge chemo-mechanical cycle. In the myosin crossbridge, we now split the unbound state into two states corresponding to myosin with ATP and ADP (states 1 and 5, respectively); we split the tightly bound state into two states corresponding to a post power stroke and a rigor like state (states 3 and 4, respectively). Transitions between states are based on the free energy differences between states, with free energy given by the free energy changes at each step in the ATP cycle plus strain potential energy in the myosin head. Rate constants and functions given in (3, 19) were used as a basis for the transition rates for the new states.

In the thick filament, forward rates depend on the free energy and the distance between the crossbridge head and the actin binding site, and there is no explicit force or velocity dependence. We define free energy as the potential energy of the two-spring crossbridge system, minus the free energy drop of each step of the ATP cycle, as measured in solution (19).

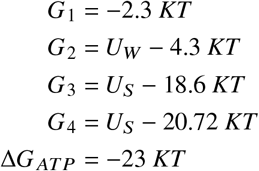

The rate equations are then calculated as:

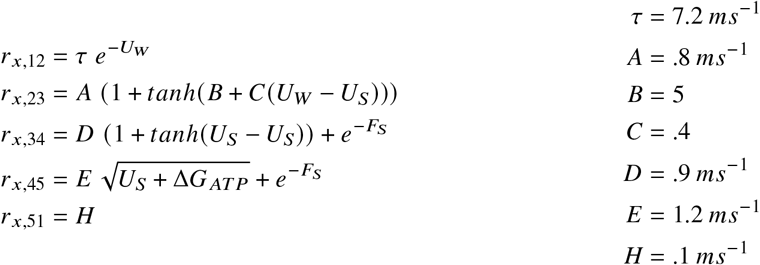

All reverse equations are defined as 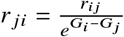. These equations are plotted in Fig. 2.

**Figure 2:**
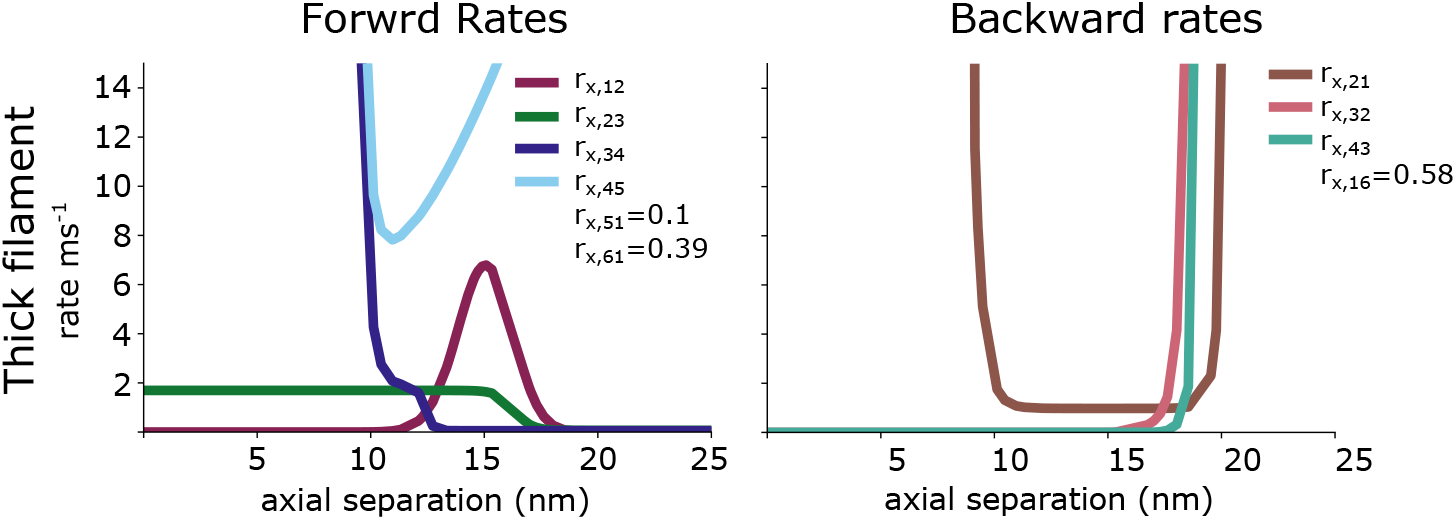
The forward and backward rate functions for the myosin binding and power stroke cycling are plotted as a function of the axial separation between a crossbridge and a binding site. The rates *r*_*x*,16_ and *r*_*x*,61_ given for a pCa of 4. Not shown are the thick filaments rates *r*_*x*,15_ and *r*_*x*,54_ as well as the thin filament rate *r*_*t*,14_ which are all 0.

We also included transitions between the disordered relaxed state (DRX, state 1) and the super relaxed state (SRX, state 6). The SRX state, also called the parked or OFF state, is a state with very low ATP turnover which allows for energy conservation in resting muscle (20, 21). The rate *r*_*x*,16_ gives the rate from the DRX to the SRX state and was chosen so that during maximal activation roughly 10% of crossbridges were in a bound state. The rate *r*_*x*,61_ gives the rate of SRX to DRX and uses the same functional form as that of (3):

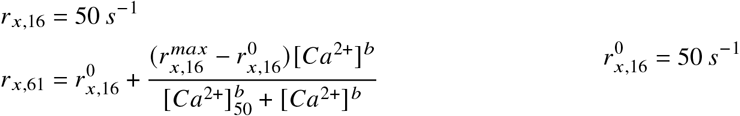

with the baseline rate 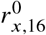 chosen so that in inactive muscle, approximately 50% of myosin heads would be in the SRX state, as found in (21, 22).

Thin filament activation is modeled as a four state system with transitions that include: *r*_*t*,12_ - Ca2+ binding to cTnC, *r*_*t*,23_ -change in the cTnC-cTnI interaction, and *r*_*t*,34_ - thin filament activation via movement of tropomyosin that allows myosin binding. Each transition in the thin filament activation pathway contains tunable forward and reverse rate constants. Reverse rates are based on the equilibrium rates below. *K*_*t*,4_ is not shown, since we define the rate *r*_*t*,14_ to be zero. Thin filament cooperative effects are modeled as a nearest neighbor effect - binding sites with a neighbor in a calcium bound state have their own forward rates multiplied by 100, as described in (14).

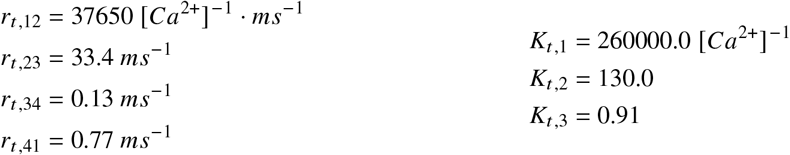

In order to simulate twitches from sarcomere models, we drive force generation and relaxation via a calcium transient, measured from isolated mouse adult cardiomyocytes. The measured Calcium transient is fit to the equation

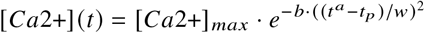

where the three modifiable parameters describe asymmetry (*a*), time to peak (*t* _*p*_) and width of the calcium trace (*w*). The constant *b* has units of 1 s(1-a) for dimensional similitude.

Another significant modification to the model involves the way we calculate the probability of state transitions. In each time step, we determine the probability of a transition for each individual myosin head and its closest neighbor actin binding site. Subsequently, we must compute the probability of transitioning from one state to another within the time-step *dt*. Previously, the probability of a transition from state *i* to state *j* was calculated simply as 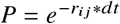. In a multi-state Markov model, however, when *r*_*ij*_ is large compared to *dt*, this leads to the sum of all probabilities being greater than one. This also does not account for the possibility of multiple transitions within a single time-step. For example, even though certain transitions may be directly impossible, we need to account for the possibility that multiple state transitions occur within a single time step. When *dt* is small relative to the rate 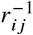 the equation is approximately correct. However, because the rate functions used have infinite walls at some strain values, indicating rapid detachment at high strains, it can become impossible to have a small enough time step.

Here, we instead rely on the matrix exponential of the rate matrix **Q**, which is the matrix which has elements *r*_*ij*_ giving the transition rate from state *i* to state *j*, with *r*_*ij*_ calculated for each possible transition based on each crossbridge’s configuration, at each time step (23, 24). The diagonal entries of **Q**, *r*_*ii*_, are chosen so that each row sums to 0, guaranteeing conservation of state transitions. Then the probability matrix **P** can be found by taking the matrix exponential: *P* = *e*^**Q**∗*dt*^. The matrix exponential is defined by 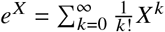, but can be efficiently approximated using Pade’s approximation (25).

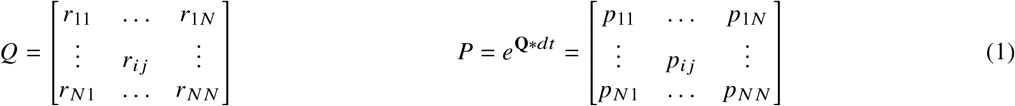

Each element *p*_*ij*_ of the matrix **P** gives the probability that if the system starts in state *i*, it will end up in state *j* after time *dt*. It also guarantees that the sum of probabilities sums to 1. So, for each crossbridge in each time step, we draw a random number from a uniform distribution from 0 to 1 and compare it with the row in **P** which corresponds to the current state of the crossbridge in order to determine the new state.

This also allows us to account for multiple transitions in a single time step as well. So for example, even though it’s impossible to go directly from state 1 to 5 in the model (*r*_*x*,15_ = 0), we can still calculate the probability of a crossbridge passing through multiple states in one time step, *p*_*x*,15_ ≠ 0. This allows us to use coarser time steps than previous instances of the model while still gaining better signal to noise. As *dt* goes to zero, **P** becomes the identity matrix and as *dt* goes to infinity, **P** will indicate the steady state distribution for the current geometric configuration and calcium concentration for each crossbridge. Because it gives the steady state only for the current geometric configuration, it cannot take into account implicit cooperative effects like compliant realignment. However, it can be used to initialize the state of the model for a given calcium concentration.

### Training data

Our training data is generated by randomly choosing multiplicative factors over the range (.1, 100) from a log uniform distribution, and every corner plot shown here shows this factor on a log scale. Each of the 9 rate functions or constants shown in Fig. 2 is multiplied by this factor, changing it from its default value that was chosen based on literature values and previous models to generate mouse cardiac twitch (2, 6, 26). Each simulated twitch was 1 second in duration, as in the experimental protocol, and consisted of 1ms time steps. The calcium transient used in our simulations was obtained from isolated mouse adult cardiomyocytes incubated in Tyrode’s Buffer with 1 *μ*M Fura-2 at room temperature in the dark for 13 minutes. After incubation, cells were moved into fresh Tyrode’s Buffer and plated on a glass coverslide at 37 °C. Cells were paced at 1 Hz and calcium transients were measured using the IonOptix Calcium and Contractility System. Because the calcium transient recorded is ratiometric, we used minimum (6) and maximum (7) pCa values previously reported in mouse trabeculae at 35 °C (27) to set the absolute magnitude of our calcium kinetics. This calcium transient was used in all simulations. We simulated each randomly selected set of rates 50 times and average them to form one data point, and we generated approximately 1.1 million rate combinations and corresponding twitches. We simulated as many rate combinations as were feasible given our computational resources to ensure that our training dataset was densely sampled enough that the probability distributions were reflective of the spatially explicit model.

While the model makes use of periodic boundary conditions, we wanted to be able to compare the magnitude of force in the spatially explicit model to experimental data. We therefore calculate the cross-sectional area of the spatially explicit model by measuring the area of the unit cell which is defined by a rhombus with vertices centered on the 4 thick filaments. This unit cell contains one thick filament (since it contains one-third of two separate thick filaments and one-sixth of two other thick filaments) and 2 thin filaments. Since the model contains 4 thick filaments and 8 thin filaments in total, we consider the cross-sectional area of the model to be 4 times the area of the unit cell. This lets us scale the output of the model, which is in *pN*, to stress, which can then be compared to experimental data, which is also reported in stress 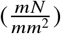.

We simulated the training dataset using Azure batch cloud computing service. Each instance of the model takes approximately 15-30 seconds to simulate, and 50 instances were averaged to form a single data point. We estimate the training dataset took approximately .25 *min* ·50 ·10^6^ /60 /24 ≈ 8000 cpu-days. We rented ‘F’ series processors from Azure cloud computing at a cost of approximately $0.015 per cpu per hour, and we were able to rent several thousand at a time, so the ‘real’ compute time was a few days. We estimate the cost of training data set to be $0.015· 1 /60 ·0.25 ·50 ·10^6^ ≈ $3000. We set 1% of the data aside as validation data, and standardized the other 99% by subtracting the global mean and dividing by the global standard deviation. These same values were used to scale validation data and experimental data for inference. Model training was performed on a Nvidia RTX 3090, and took approximately 24 hours. After the model is trained, generating a probability distributions corresponding to a target twitch takes approximately a minute. While generation of the training data can be computationally intensive, it has the advantage of only needing to be trained once, and parameter inference of further targets requires only minimal computational effort.

### Conditional Variational Auto-encoder (CVAE)

Auto-encoders are a type of machine learning tool which can be used for both dimensionality reduction and generative purposes. In general, they consist of encoder and decoder layers, where the dimensionality of the encoding layer’s output (often referred to as the latent space, which we denote as *z*) is much smaller than the input, which allows them to be used for dimensionality reduction. Variational auto-encoders are a particular type of auto-encoder in which the output of the encoder is designed to describe a probability distribution within the latent space rather than a single point. A single point drawn from that space is then used as the input to decoder. We also can condition the input to the auto-encoder on an observation, in our case, the twitch force 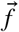.

The design of this specific Conditional Variational Auto-encoder (CVAE) follows that described in (8). It consists of three sub-networks: two encoders, *Q*_1_ and *R*_1_, and the decoder *R*_2_ (Fig. 3). Both encoders and the decoder have the same general architecture, consisting of a convolutional ResNet followed by a series of fully connected layers with only the width of the first fully connected layer and size of the output differing between each of the sub-networks. The convolutional layers are shared by each encoder and decoder, meaning they use the same weights.

**Figure 3:**
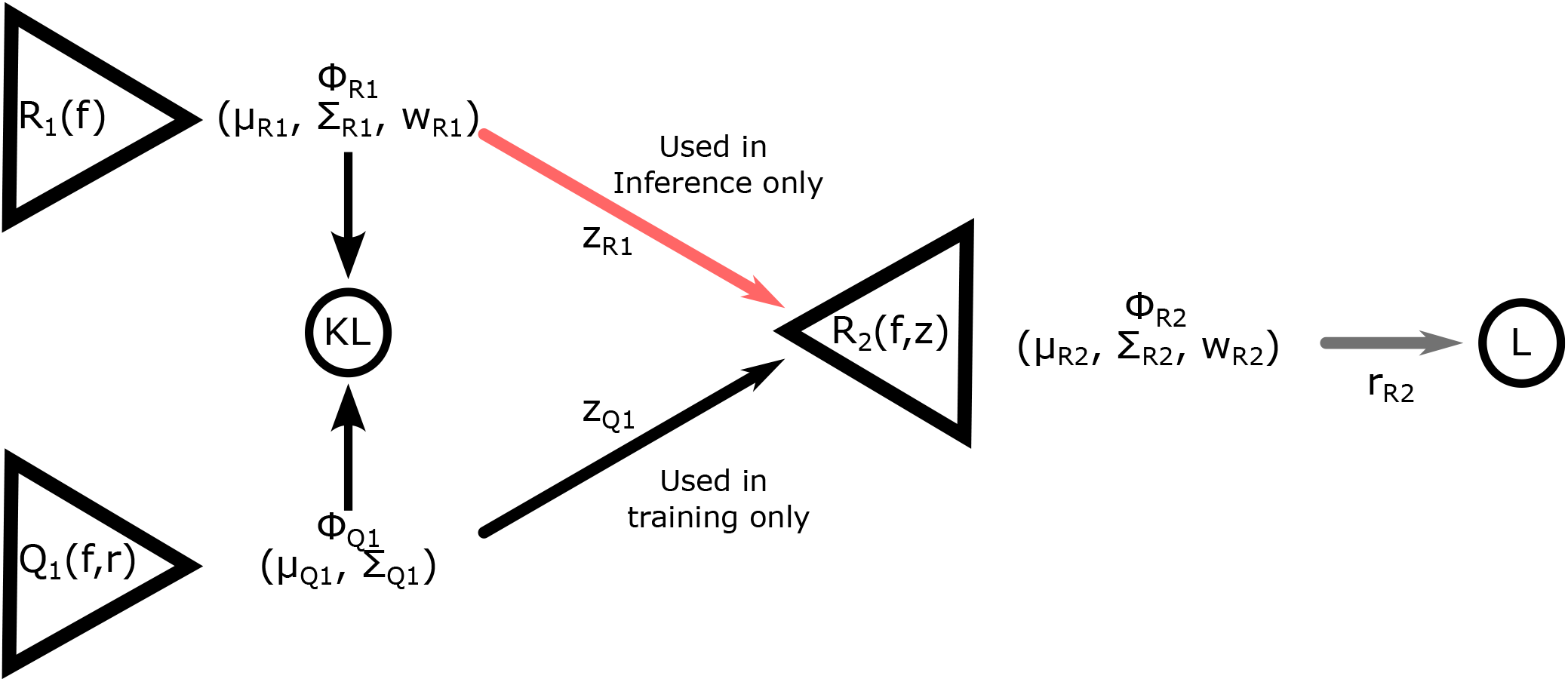
The encoding and decoding networks *Q*_1_, *R*_1_, and *R*_2_ are each composed of the same shared deep convolutional layers. The loss function -L+KL is composed of the log probability of the true rate at for the estimated probability distribution function (L), and the Kullback-Liebler divergence (KL) between the probability distributions from the *Q*_1_ and *R*_1_ networks (KL). 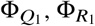, and 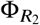 are the probability distributions generated by the separate networks, and *μ*, Σ, and *w* are the means, variances, and weights of the components of each of the Gaussian Mixture distributions. The points 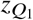 and 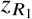 are random points drawn from their respective distributions over the latent space, and 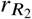 is a random rate drawn from its distribution. This architecture follows that found in (8).

In the encoder *Q*_1_, after the convolutional layers, the convolved twitch is concatenated with the true simulation rate parameters *r* before being passed to the fully connected layers which outputs the means and log-variances describing a unimodal *n*_*z*_-dimensional Gaussian distribution 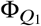. In the encoder *R*_1_ only the convolved twitch is passed to the fully connected layers, which outputs the means, (log) variances, and (log) weights describing an *n*_*z*_-dimensional Gaussian mixture model with *m* components, 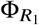. The KL divergence between these two distributions forms part of the loss function.

In the decoder *R*_2_, after performing the same convolutional operation on the force time-series, a point is drawn from either the distribution 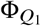 or 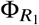, depending on if we are in training or inference mode. During training, the point is drawn from the unimodal 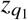, which was given the true rate parameters as part of its input, while during inference the mixture distribution 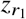 is sampled. In either case, this *n*_*z*_-dimensional point is concatenated to the convolved time-series and passed to the decoder *R*_2_, which will output the parameters of an *n*_*r*_ -dimensional, *m*-component Gaussian mixture model, 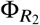.

The full derivation of the loss function is given in (8, 28). In short, we start with the cross-entropy loss between the ‘true’ distribution *p* (*r* | *f*), and our estimate of the distribution generated by the CVAE, *R*(*r* | *f*):

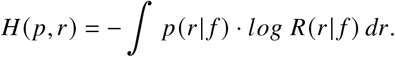

Since the the distribution *r* (*r*| *f*) is actually the product of the distributions of the encoder/decoder pair *R*_1_ and *R*_2_ marginalized over the latent variable z, we can write it as

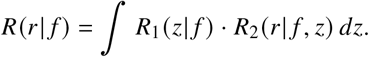

During training, however, we use the simpler encoder/decoder pair *Q*_1_ and *R*_2_, also called the recognition function

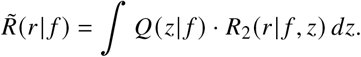

By finding the Kullback-Liebler divergence between these two networks, and substituting the difference into the equation for *H*(*p*|*r*), and minimizing the expectation of H over our observed data f, we can derive that the difference between the ‘true’ distribution *p* (*r* | *f*) and the CVAE distribution *r* (*r* | *f*) can be written as

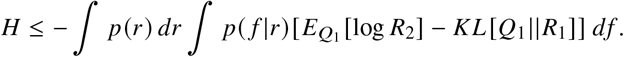

We approximate this integral as

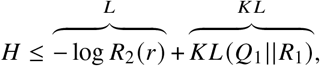

The loss function consists of two values, the log-probability *L* of the distribution *R*_2_ at the location of the true rate parameters *r*, and the Kullback-Leibler (KL) divergence between the distributions *Q*_1_ and *R*1. The value − *L* +*K L* is minimized. Similar to (8), in practice we minimize the value − *L* +*α* ∗*K L*, where *α* is 0 for the first 30 epochs, and then linearly increases for the next 60 epochs to one. This is done because the CVAE network can become stuck in local minima where the KL term is at its minimum of 0, which prevents optimization of the L term.

The function we are interested in, *P* (*r* |*f)*, is not in a closed form, but is modeled by the networks *R*_1_ and *R*_2_ together. The network *R*_1_ yields a distribution conditioned on the target twitch, from which we draw a single point 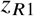 used to condition the distribution 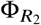 from which we draw a single rate factor 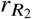. The network must then be sampled many times to obtain the distribution *P* (*r* |*f)*. To visualize *P* (*r* |*f)*, we sample the network 5000 times to obtain a histogram of rate factor combinations. We use the ‘pairplot’ function from Seaborn (29), which takes the histogram and fits and plots a distribution using a kernel density estimator.

Besides the convergence of the loss function, we evaluate the model’s performance using the probability-probability plot, as in (8), to ensure that the probability density estimate we find is representative of the actual probability density. For an ideal estimator, we should expect that when we estimate the probability that the true parameter lies with a certain volume of the rate space to be X%, we should expect the true parameter to be in that volume X% of the time. Thus, a plot of the fraction of times the true rate was found within the credible region for our validation dataset provides a measure of the model’s performance. The diagonal of the plot would represent a perfect estimate.Values above the diagonal indicate under-confidence, and values below indicate over-confidence in the estimation. We show this probability-probability plot in Fig. 4, with each colored line representing one of the nine rates and the black dashed line indicating the ideal case. These were constructed by first finding the cumulative 1-D marginal distribution for each rate for each data point (twitch) in our validation set. We then plotted the fraction of posterior samples less than the true rate parameter. Our testing set contained approximately 11000 samples, or 1% of our total simulated dataset, which is adequate given the large training dataset and provides low error in the probability estimate (test sets above 1000 converge to the same low error: Supplemental Figure 5).

**Figure 4:**
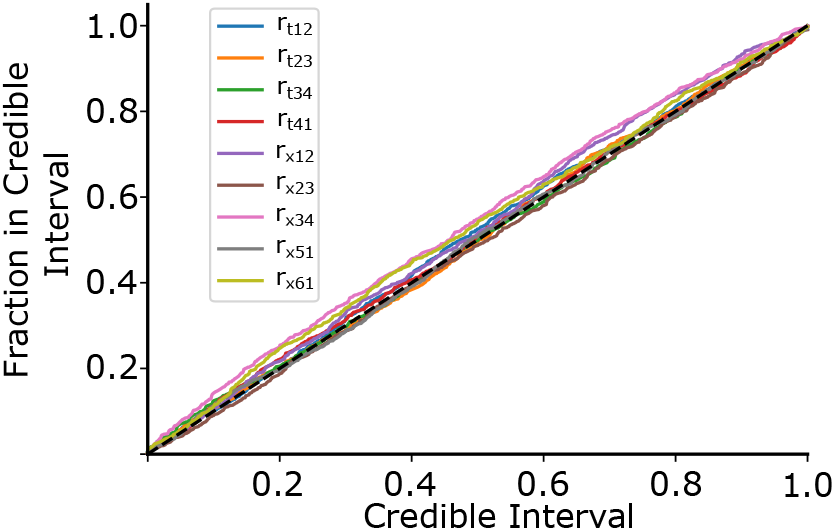
The Probability-Probability plot shows how well our estimated probability distribution describes the actual probability distribution across our validation dataset. The x-axis shows the estimated probability the rate is contained in a certain volume of the rate space, and the y-axis indicates the actual fraction of times the rate was contained in that volume. For an ideal predictor, the true parameter should be in the X% credible region X% of the time, indicated by the black dashed line. Values above the diagonal indicate under-confidence, and values below indicate over-confidence.

**Figure 5:**
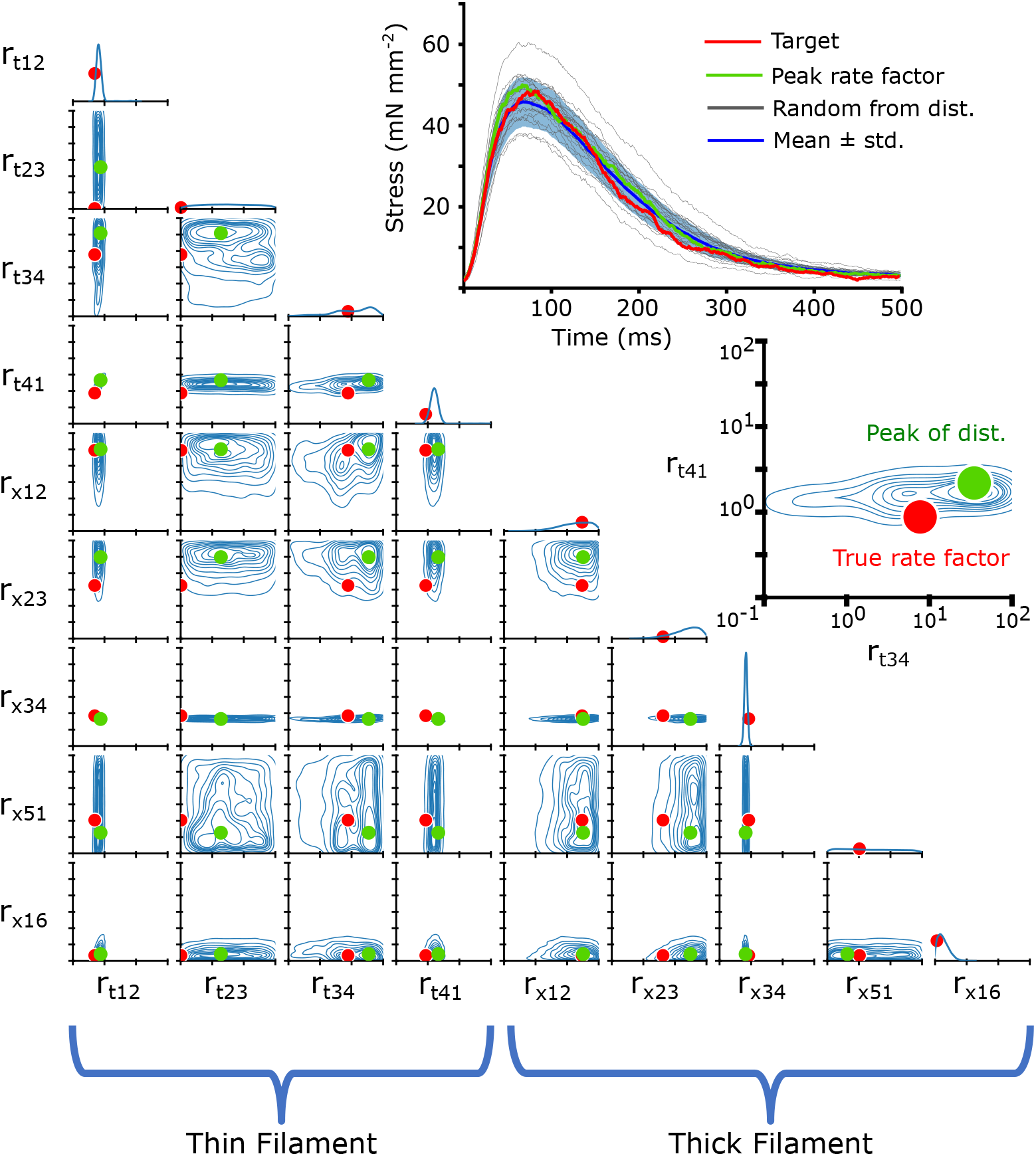
The 2-D joint probability distributions for the 9 rates we chose to vary in our simulations are shown as contour plots that indicate the probability density over the rate space. The 1-D marginalized probability distribution of every rate factor is plotted along the diagonal. In each joint probability distribution, solid red dots indicate the true rate factor which the target twitch had been generated with. The target twitch, drawn from the simulated validation subset, is shown in the upper right in red. A second inset shows a blowup of the joint probability distribution for *r*_*t*,12_ and *r*_*t*,34_, with the true rate factor indicated as a solid red dot. We also show the peak of the distribution as a solid green dot, and the resulting twitch simulated with that rate factor combination is shown in the upper right in green. Also shown are 20 twitches (black) and their mean ± standard deviation (blue) which were simulated with rate factors drawn randomly from the distribution. The rate factors are multiplied by the base rates, plotted in Fig. 2. Note that the rate factors are plotted on a log scale.

## RESULTS

### Parameter Inference of a Simulated Twitch

After training the CVAE network on a simulated training dataset, as described in the Methods section, we can illustrate the probability density function for all of the nine parameters that underlie a specified twitch. By examining only the time series of the twitch, and comparing to the known rate factors which led to the simulated twitch, we can estimate the accuracy of its parameter inference on unseen data.

In order to visualize the posterior distribution 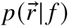, we show the resulting multidimensional distribution as a corner plot. In the corner plot, every 2-D projection of the high dimensional distribution is shown. This allows us to see every 2-D marginalized probability distribution. Also, along the diagonal, we can show the 1-D marginalized probability distribution of every rate factor. The axes indicate the rate factors that were multiplied by the base rates, and are shown on a log scale from 10^−1^ to 10^2^. An example simulated twitch and inferred distributions are shown in Fig. 5.

Corner plots allow us to visualize the covariance between each 2-D pairwise parameter combination. For example, we can see in Fig. 5 the covariance between the rate factors *r*_*t*,12_ and *r*_*t*,41_, indicating that changes in one parameter can potentially be partially compensated for by changes in the other. We can also interpret the uncertainty as a kind of parameter sensitivity analysis, indicating, for example, that the rate *r*_*x*,34_ must be tightly constrained for this particular simulated twitch.

### Control and I61Q cTnC twitches from mouse trabeculae

We wanted to see how our CVAE would perform on inferring rates in real muscle, so we collected twitch data from mouse trabeculae. We used control and I61Q cTnC mice, which we have previously shown to model genetic cardiomyopathy with hypocontractility at cellular and organ level (11, 12). The trabeculae were held isometrically and stimulated at 1 Hz while stress was recorded (see Methods for further details). We recorded 10 twitches from 8 different muscles for both control and I61Q cTnC transgenic mice. We averaged the twitches together for each type and show the mean ± standard deviation in Fig. 6. Consistent with previous findings (11, 12), we found that the I61Q cTnC variant resulted in reduced stress relative to the control twitch by approximately half. We also used previously published data on mouse trabeculae treated with Danicamtiv, as well as their accompanying control data, and inferred rate distributions for those twitches as well. Table 1 shows mean and standard deviation of the peak stress, as well as peak, rise, and fall timings of stress for all conditions.

**Table 1:** Stress and timing differences - Control, I61Q cTnC, and Danicamtiv and its control mice trabeculae. Danicamtiv - and its matched control data - first published in (12). We define *t* _*peak*_ as time to peak stress, *t*_50,*rising*_ as time to 50% maximum stress and *t*_50, *falling*_ as time to 50% relaxation, relative to the start of activation.

|  | Control | I61Q cTnC | Control | Danicamtiv |
| --- | --- | --- | --- | --- |
| Peak Stress | $59.8 \pm 13.0 \frac{mN}{mm^2}$ | $28.9 \pm 9.3 \frac{mN}{mm^2}$ | $25.1 \pm 14.3 \frac{mN}{mm^2}$ | $40.1 \pm 16.0 \frac{mN}{mm^2}$ |
| $t_{peak}$ | $98.7 \pm 18.6$ ms | $106.2 \pm 9.6$ ms | $119.5 \pm 4.1$ ms | $140.7 \pm 11.4$ ms |
| $t_{50,rising}$ | $45.1 \pm 5.8$ ms | $48.2 \pm 3.1$ ms | $59.5 \pm 2.8$ ms | $68.5 \pm 7.1$ ms |
| $t_{50,falling}$ | $166.8 \pm 33.9$ ms | $173.5 \pm 15.0$ ms | $185.8 \pm 8.5$ ms | $224.8 \pm 17.5$ ms |

**Figure 6:**
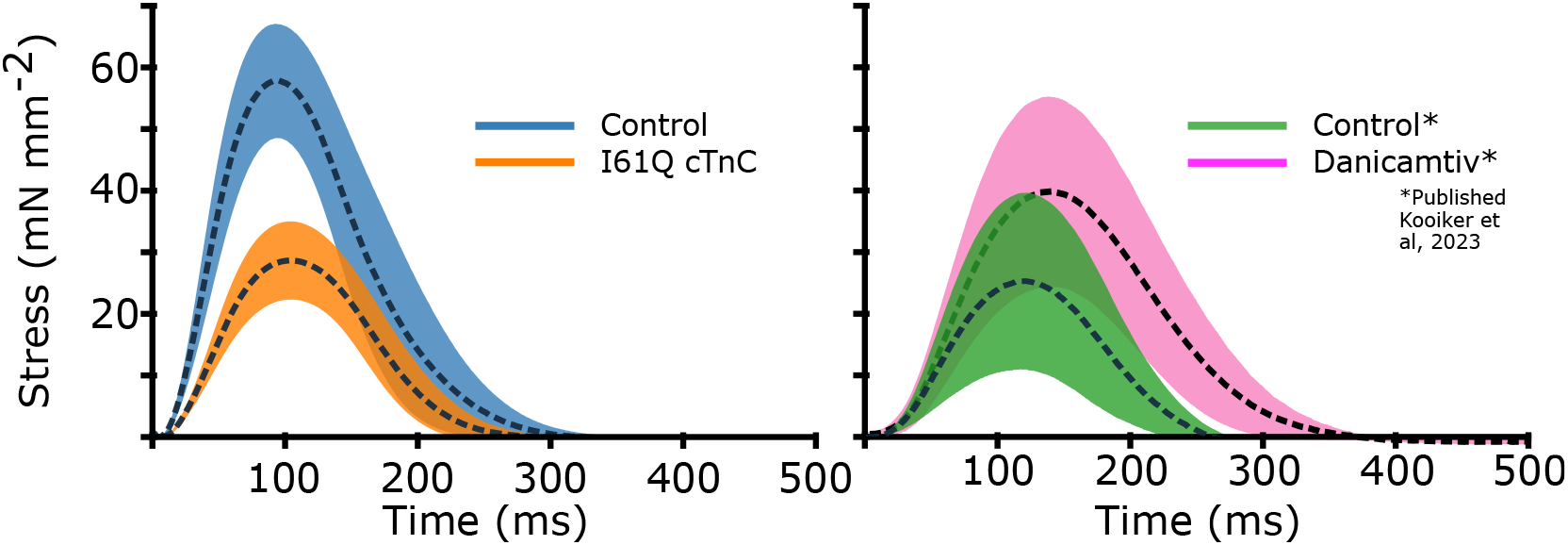
Here we show the experimental isometric twitch stress for the different conditions used in the CVAE. First, we show mouse cardiac trabeculae from control (blue) and mice with the I61Q cTnC variant (orange). We also show control trabecula (green) and trabecula treated with the myosin activator Danicamtiv (pink), previously published in (12). We show the control datasets taken contemporaneously for both I61Q cTnC and the Danicamtiv, since a different experimental apparatus and protocol was used. For each group, the overall average (dashed black line) is shown along with the 95% confidence of the mean intervals (shaded region). The I61Q cTnC groups consisted of 8 individuals, while the Danicamtiv groups consisted of 4 individuals.

### Parameter Inference of Mouse Cardiac Twitches

Using the mean experimental twitches for control and I61Q cTnC across individuals (Fig. 6), we next used the trained CVAE to infer the set of rates which were most likely to lead to a simulated twitch which was similar to the experimental twitch. Fig 7 shows the predicted probability distributions for the mean experimental twitches of both the control (blue) and I61Q cTnC (orange), with the most probable values indicated. One notable result from the probability distribution plots is that rate constants for the calcium thin filament on and off rates (*r*_*t*,12_ and *r*_*t*,41_) are among the most divergent between the two twitches, which is consistent with the known experimental data of the I61Q cTnC variant (30). To further demonstrate utility, we applied the CAVE method to our previously published cardiac mouse twitches ((12)) that were collected in presence and absence of the myosin activator Danicamtiv and show a different probability distribution compared to the I61Q cTnC twitches. The CVAE method correctly identified the two underlying mechanisms of Danicamtiv, which are recruiting more myosin motors into the “On” state (decreased *r*_*x*,16_) and slower ADP release(decreased *r*_*x*,34_). The results demonstrate the strength of the CVAE as it was not provided any information about the underlying mechanism of the abnormal twitches.

**Figure 7:**
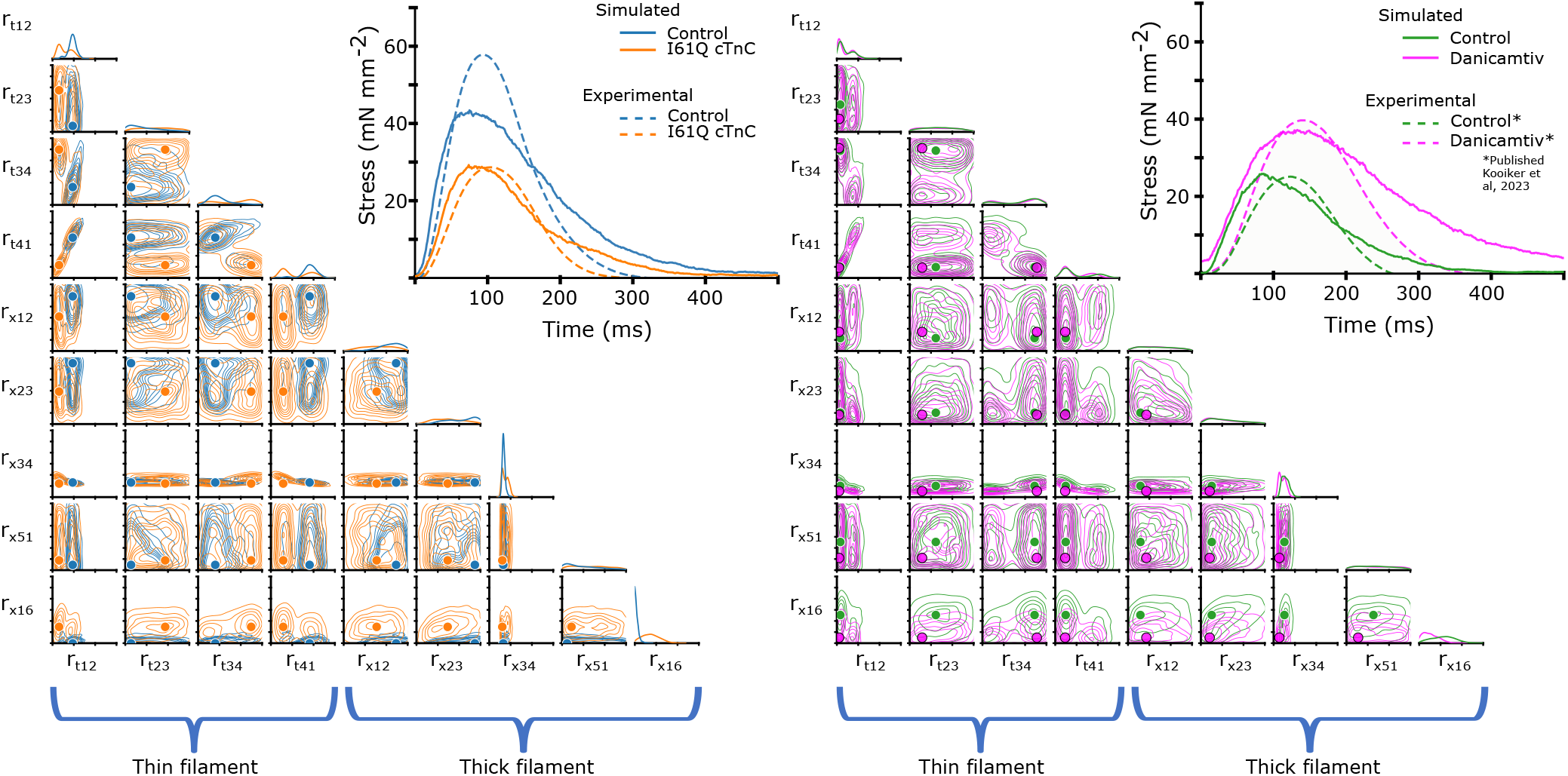
As in Fig. 5, we show the 2-D joint probability distributions as contour plots that indicate the probability density over the rate factor space. Here, however, our target twitches were the mean experimental data shown in Fig. 6 rather than simulated data, and dots indicate the peak of the probability distribution, i.e. the most probable value. In the left panel, we show the experimental twitches as well as the twitch simulated using rate parameters corresponding to the peak of the respective distribution. In the right panel, we show the same using data previously published in (12). As before, the rate factors are multiplied by the base rates, plotted in Fig. 2. Note that the rate factors are plotted on a log scale.

Next, we wanted to use the probability distributions plots to determine how well we can recapture the experimental twitches. Using the most probable (peak) value of the estimated distribution, we simulated a twitch using our spatially explicit model and plotted the resulting simulations along with the experimental twitches (Fig 7, upper right). In general, the simulated twitches using the rate parameters estimated by the CVAE follow the rising and falling characteristics of the simulated twitches, but do not achieve the same peak force. This is particularly noticeable in the control from the I61Q cTnC group and the Danicamtiv treated twitches.

### Thick filament intervention to correct thin filament Ca^2+^ deficiency

In the parameter inference above, we allowed the CVAE to vary all nine rates in order to best fit the experimental data in both control and I61Q cTnC variants. However, we know from published experimental work that the primary mechanism by which the I61Q cTnC variant decreases muscle performance is by reduced Ca^2+^ binding affinity (30). We decided therefore to start from the control parameters inferred in Fig. 7 (blue) and then selectively alter the two parameters affecting Ca^2+^ affinity, *r*_*t*,12_ and *r*_*t*,41_, to match that of the inferred I61Q cTnC twitch (orange). This hybrid twitch, shown in Fig. 8, therefore combines the inference of the CVAE with experimental knowledge to provide an estimate of an I61Q cTnC twitch. The I61Q cTnC simulation with this hybrid rate factor combination results in similar kinetics as the control simulations, allowing us to evaluate potential non-thin filament therapeutic targets to recover the twitch phenotype.

**Figure 8:**
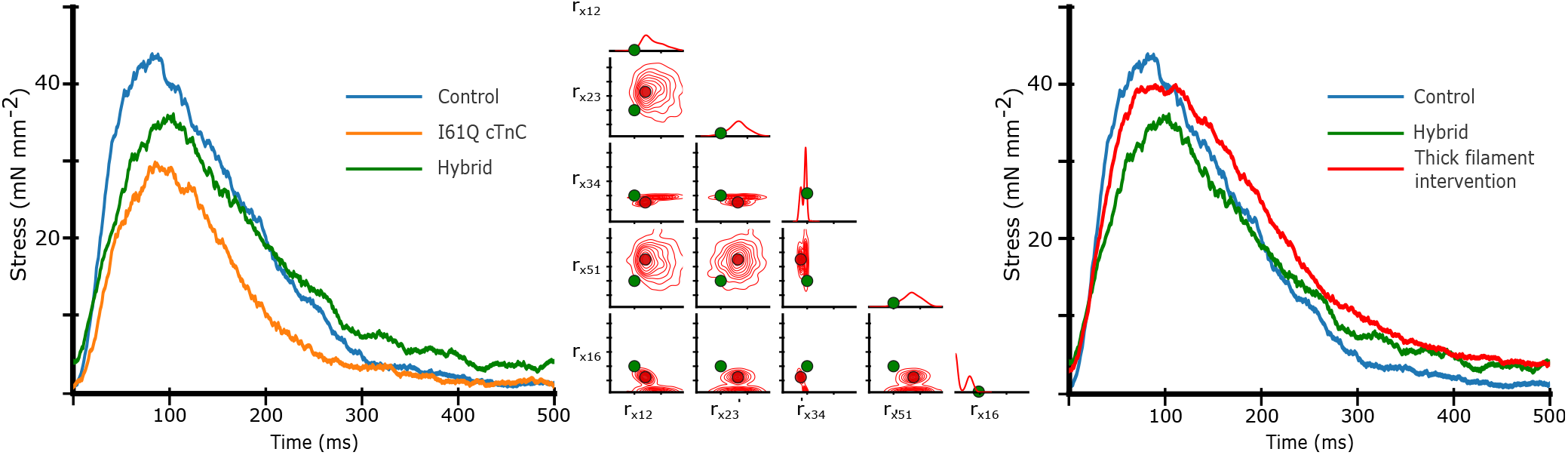
Left) Isometric twitch stress is plotted against time. Dotted lines indicate the experimental data, and solid lines indicate simulated data. The blue and and orange simulations are performed with the most probable rates identi from the distributions shown in figure 7. The ‘hybrid’ twitch (green) combines rates from those two inferences: the *r*_*t*,12_ and *r*_*t*,14_ rates from I61Q, and the control rates for the rest. Middle) Starting with the ‘hybrid’ twitch and using the Control twitch as a target, we trained a separate CVAE on only the thick filament rates in order to estimate an ‘intervention’ in the thick filament, i.e. changes we could make which would restore function. For this set of simulations we centered the training set around the ‘Hybrid’ rate factors, represented by green dots.Here, we show the resulting distribution over the thick filament rates using the control twitch as a target. As in Fig. 7, solid red dots indicate the most probable rate factor, and rate factors are plotted on a log scale. Right) We show the resulting simulated twitch corresponding to the peak of the distribution, which is our best estimate for a thick filament intervention. In short, we started with the control twitch (blue), reduced *r*_*t*,12_ and *r*_*t*,14_ to estimate an I61Q twitch (green), then estimate what change would be necessary to restore function, i.e. give a twitch most similar to the control (red).

Starting from the hybrid twitch we simulated in Fig. 8, we asked what intervention(s)(in simulation) could potentially correct for the alteration in thin filament Ca^2+^ sensitivity we introduced. To accomplish this, we generated a second data set in which we fixed the rate factors in the thin filament that were used in the hybrid twitch (Fig. 8) and allowed the rates in the thick filament to vary as before, from .1 to 100 over a log uniform scale. We then trained a second CVAE network to predict the probability distribution of rates over this reduced space. Once the CVAE had been trained sufficiently (as determined by the training curve in conjunction with a prob-prob, as in Fig. 4), we used the original control twitch as input and generated a probability distribution over the thick filament rates. We then chose the rate factors corresponding to the most probable (peak) value of the probability distribution as input to the spatially explicit model. The probability distribution, most probable rate factor, and resulting twitch are shown in Fig. 8. We found that the estimated intervention approximately restored the peak twitch force, but that the relaxation time had been increased in comparison to the original control simulation. These findings are similar to our published work using the myosin activator Danicamtiv to recover the abnormal twitch of the I61Q cTnC mice (12).

## DISCUSSION

In this study, we wanted to combine Bayesian inference methods with a spatially explicit muscle model in order to predict combination of transition rates that can reproduce the isometric twitches from healthy and diseases mouse cardiac trabeculae. By re-framing this inverse problem using Bayes’ law, we can infer a probability distribution over the rate space - rather than single rate values - which may generate simulated twitches matching experimental data. Prior studies have shown that MCMC methods for parameter estimation in inverse problems can be more computationally efficient than the CVAE approach (8). Such alternate approaches would be potentially useful in the case that only a single target twitch.But methods such as the CVAE can be more efficient when there are a broad range of target data of interest. As we mentioned above, while the generation of the training data can be computationally intensive, it has the advantage of only needing to be trained once, and parameter inference of further targets requires only minimal computational effort. We hope that by continuing with such methods, it may be easier for a broad range of researchers - who may all be interested in different small molecule modulators or different genetic variants - to get quick estimates for a range of parameters, and generate hypothesis on which steps of the crossbridge cycle may be effected most by a particular intervention. By finding such distributions for both control as well as different genetic variants in mouse cardiac trabeculae, we hope to further be able to develop a method to infer how changes in some rate factors can ‘undo’ changes made in another.

We began by updating our previously published computational model of the sarcomere to include additional states in order to more easily compare the spatially explicit muscle model to experimental results. The new model was used to generate a large data set of cardiac twitches in which 9 of the model parameters were randomly varied. We then used Bayesian parameter estimation with a Conditional Variational Autoencoder (CVAE) to generate probability density plots of parameter(s) that result in a particular cardiac twitch.

Four key results emerge from our study. First, changes to probability calculations allow for more accurate simulations with fewer times-steps and less variations. This will have implications for others who use similar stochastic models of muscle contraction. Second, variational inference enables accurate parameter estimation for high dimensional data such as muscle twitch. Third, machine learning tools such as conditional variational autoencoders that are trained on simulated data, can be applied to experimental twitch data. We are able to predict the relative importance of model parameters that can produce particular normal or diseased twitches. Fourth, we are able to use our newly trained tools to predict possible ways to recover abnormal cardiac twitches.

Before we expand on the results, it is important to highlight the limitations of this study. First, the spatially explicit model only captures multiscale interactions at the half-sarcomere level while in whole muscle, interactions between heterogeneous collections of half sarcomeres as well as structures outside the sarcomere may have significant effects on whole muscle function (5, 31–33). A whole muscle is not simply a half sarcomere scaled up as we have done here. Despite this, is has been shown that isometric twitch stress - simulated and experimental - can be predictive of underlying biophysical perturbations (34). Another challenge is the vast number of parameters which could potentially be explored. Our spatially explicit model contains parameters that kinetic, mechanical, and geometric aspects of the sarcomere. Kinetic parameters include the transition between states in the actin binding states and myosin crossbridges, such as the nine rate factors we modified in this paper. However, those nine are only a subset of the overall rate space that could be affected by genetic variants. Mechanical parameters represent the elastic coupling between neighboring myosin heads and binding sites, as well as the mechanical properties of the myosin heads themselves, and changes in that coupling can lead to very different sarcomeric function (2). Geometric parameters represent the spatial distribution of myosin heads including their orientation, number relative to thin filaments and radial spacing, all of which can depend on species (35, 36). Because the model is spatially explicit, any sub-population, or even multiple sub-populations, can be given different characteristics, leading to an exceedingly large parameter space. And, because of the spatially explicit nature of the underlying model, each value can be varied. Spatial variation in parameters can be useful, since the penetrance of a genetic variant may not be 100%, but rather only a sub-set of proteins may be affected, and their distribution may be uniformly random or spatially heterogeneous (34). Another limitation of our study is that we used the same calcium transient for all simulations. We and others have shown that alterations in myofilament function results in changes in calcium dynamics (11, 37, 38). We elected to only look at sarcomeric parameters, to keep the parameter space more manageable.

### Changes to probability calculations allow for more accurate calculations

One significant modification to the spatially explicit muscle model is the implementation of the matrix exponential in obtaining the per time step probability of transition for states. Previous work with our model used fewer states and were isometric, in which the approximate probability calculations were sufficient. Thus our model, and other similar Monte-Carlo simulations, simply used the rate and simulation time step to compute a transition probability (7, 39). However, with the addition of states this approximation should become less accurate. For example an unbound crossbridge in state 1 could transition into one of three states. Using our previous probability calculation, each competing state would require a smaller and smaller time step for an accurate approximation of the probability of transition, increasing computational cost. Using Eq. 1, since the probability is guaranteed to be conserved regardless of time-step size, we can use coarser time steps. This speeds calculations not only by having to simulate fewer time-steps per twitch, but because variation is reduced, we can average fewer twitches together to form each data point in the training dataset (Fig 3). Due to these changes in the probability estimation, we expect that techniques with higher frequency oscillations and amplitudes, such as Nyquist measurements in (40), should be feasible.

### Variational inference allows for approximation of high dimensional inverse problems

In general, high dimensional inverse problems are difficult for a number of reasons, namely that there is often no unique solution for a particular observation, but rather many solutions which lie on a manifold in a high dimensional space. In other words if we observe some data 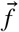, we can expect for a theoretical model, there will be many parameters 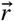 which can generate that data (41).This problem is further complicated by the addition of noise in a model. A common way to deal with this under-determined nature of the problem is by adding regularization terms to a cost function. By adding additional constraints, one can limit the number of potential solutions (42). By reformulating the problem in terms of Bayes Theorem, we can model the solution as a random variable and guess its probability distribution instead of treating it as a single value.

We can then interpret the probability distribution as indicating the relative importance of one parameter compared to another. The probability density plots indicate a measure of the covariance of parameters, or a kind of sensitivity analysis, with broad uncertainties implying changing that particular parameter will have a relatively small effect on the resultant simulated twitch compared to parameters with low uncertainty. Even in small training datasets, it can become apparent if a particular parameter is especially dominant. However, ensuring the training set is dense enough is necessary to ensure uncertainties are reflective of the ‘true’ covariance of the different parameters. For the density of training data to remain constant as the number of parameters increase, the number of training data points must increase exponentially with the number of dimensions (parameters). We chose the rate factors in this paper due to a combination of being important from a computational standpoint, as well as biologically relevant due to the known effects of specific genetic variants (11, 30) and small molecule modulators (12, 43).

### Combination of Simulations and ML methods can illuminate underlying mechanisms of muscle diseases and ways to recover function

Machine learning methods such as variational autoencoders are increasingly applied in a variety of biological settings to identify cell types that contribute to disease states (44), design novel protein variants (45) or recombinases (46). Recently, autoencoders have been applied to multiplexed immunofluorescence images to understand how alterations in different processes can alter subcellular organization (47). For the first time, we apply CVAE to cardiac twitch simulated and experimental data.

As seen in Figures 7, this method can produce probability distributions which can inform us of the relative importance of certain rates to a particular target twitch. This method can then aid in generating hypotheses about underlying cardiac sarcomere biology. For example, in Figure 5, there is very little variation in the predicted *r*_*x*,34_ for the single twitch shown. This is consistent with the fact that ADP release is the rate-limiting step in the crossbridge cycle and cardiac muscle relaxation kinetics.

The traditional way of studying sarcomeric pathology is to start with biochemical or biophysical assays often from recombinant proteins and then scale up to cellular, tissue or animal models. Based on the mechanism of action, potential treatment strategies can be tested *in vitro* and then *in vivo*. This approach is informative for single variants but lacks the resolution to screen many variants at once. Here we apply novel ML tools to muscle twitch data to infer underlying mechanism of action. As seen in Figure 7, there are many possible set of parameter combinations that can perform reasonably well in generating a twitch that resembles an average control or I61Q cTnC. For many of the thick filament parameters, there is significant overlap between the rates for these two conditions. However, for most of the thin filament parameters, there is very little overlap between the possible rates. As seen in Figure 7, there is minimal overlap between the *r*_*t*,12_ and *r*_*t*,41_parameter space between control and I61Q cTnC. These findings are consistent with the known mechanism of I61Q cTnC variant, which reduces Ca^2+^ binding affinity and cTnC-cTnI interaction (30). This demonstrates the power of this technique as the trained model did not know anything about the underlying mechanism of the diseased twitch.In order to demonstrate utility in a range of conditions, we applied CVAE to our previously published mouse cardiac twitches treated with the myosin activator Danicamtiv (12). As seen in the left panel of Figure 7, we were able to show changes on the thick filament parameters, namely decreased *r*_*x*,34_ and decreased *r*_*x*,16_, which corresponds to our previously published on the primary mechanism of Danicamtiv (12).

To help in determining which rates account for the most difference between two twitches, we used the Kolmogorov–Smirnov (KS) test. The KS statistic informs about the maximum difference between the two cumulative distribution functions (CDFs) and the sign indicates which CDF is larger with a negative sign indicating that the value for the control CDF was smaller. In case of the control vs. I61Q cTnC, the KS statistic for *r*_*t*,12_ and *r*_*t*,41_ distributions are higher than most other parameters, consistent with the primary mechanism of action of the variant. One exception, however, is the rate *r*_*x*,16_. The results are even more clear when comparing control and Danicamtiv twitches, as the KS statistic for the rates *r*_*x*,34_ and *r*_*x*,16_ are larger than the other rates, consistent with the hypothesis that changes in those rates account for most of the difference in those twitches.

Another use of of ML methods in cardiac biology is in predicting potential therapeutic targets. While there are no examples of this approach in sarcomere biology, a recent study used ML methods to identify a small molecule that corrected gene network dysregulation in NOTCH1-haploinsufficient hiPSC-derived endothelial cells (48). As seen in Figure 8, we generated a twitch with the best predicted thin filament parameters that fit the I61Q cTnC data (orange trace). We then generated a new smaller data set where only the thick filament parameters are varied in order to tell us about potential thick filament-based interventions to correct the abnormal twitch. Two effective strategies seem to be reducing ADP release rate (decreasing *r*_*x*,34_) and increasing the DRX myosin population (decreasing *r*_*x*,16_). As discussed earlier, these two rate constants are affected by the myosin activator Danicamtiv, which we have previously shown to correct the hypocontractility seen in the I61Q cTnC mouse model (12). While the peak force is clearly recovered by using this type of approach, the relaxation is prolonged, which we also saw in our experimental treatments with Danicamtiv.

## AUTHOR CONTRIBUTIONS

TT, FMH, TD designed research and analyzed data. TT performed simulations and wrote ML model. KK collected the experimental data. TT, FMH, TD, KK, JD wrote and edited the manuscript.

## DATA AVAILABILITY

The spatially explicit model used in this article can be found at https://github.com/travistune3/multifil_five_state. The training and testing datasets are available from Dryad at DOI: 10.5061/dryad.d51c5b0bj. The CVAE model is available at https://github.com/travistune3/CVAE.

## ACKNOWLEDGMENTS

This work was supported by NIH grants R01HL157169 (FMH), R01HL142624 (JD) and an American Heart Association Collaborative Sciences Award (FMH, JD and TD). This research used resources of the University of Washington Center for Translational Muscle Research, supported by the NIH National Institute of Arthritis and Musculoskeletal and Skin Diseases under Award Number P30AR074990 (TD and JD). Cloud computing credits for this work were supported by the University of Washington eScience Institute in partnership with Microsoft Azure.

## SUPPLEMENTARY MATERIAL

**Supplementary Figure 1:**
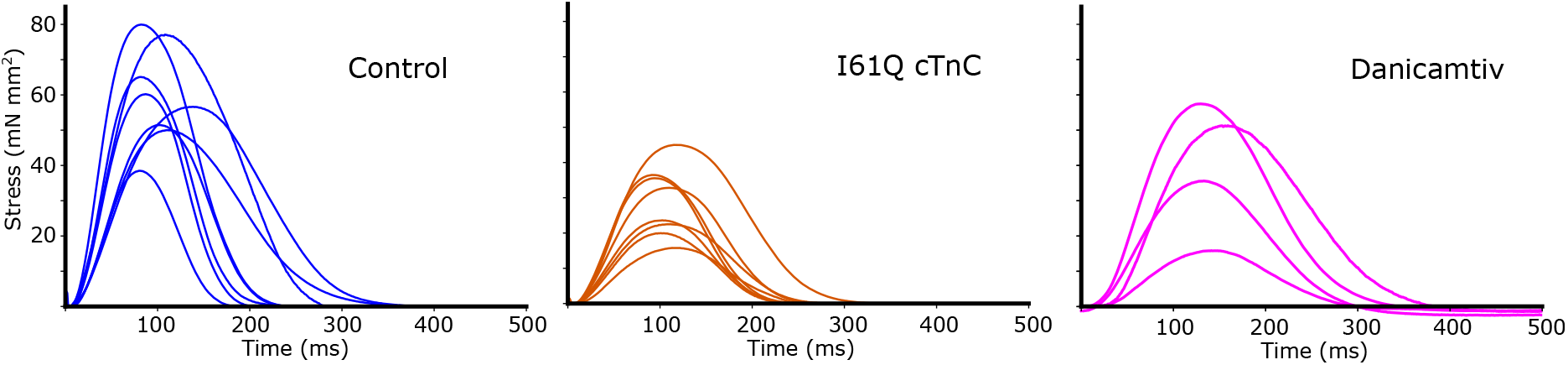
Here we show twitch force in mice cardiac trabeculae from both control (blue) and mice with the I61Q cTnC variant (orange). Black dotted lines indicate the twitches from 8 different individual mice, each of which is an average of 10 twitches. The overall average is shown with 95% confidence of the mean intervals.

### Mouse Cardiac Trabeculae Isometric Twitches

Here we show the individual traces of all 8 individual muscles for both control and I61Q cTnC mouse cardiac trabeculae, as well as the 4 twitches which make up the danicamtiv dataset. Each individual trace is composed of 10 twitches which are averaged (Fig. 1). Danicamtiv data was first published in (12).

### Mean and Variance in exp data and model predictions

Because experimental data often has high variance between individual measurements, we wanted to see how the CVAE performed when shown different instances of the same type of twitch. Here we inferred the set of rates associated with the overall average twitch for a control muscle and plotted the resulting rate distribution. Then we inferred the rates for each of the 8 individual muscles separately, and then found the average rate distribution across those 8 distributions. The results show, in general, that the average of the separate predictions has a larger variance, but that the two methods produce the same means (Fig. 2).

**Supplementary Figure 2:**
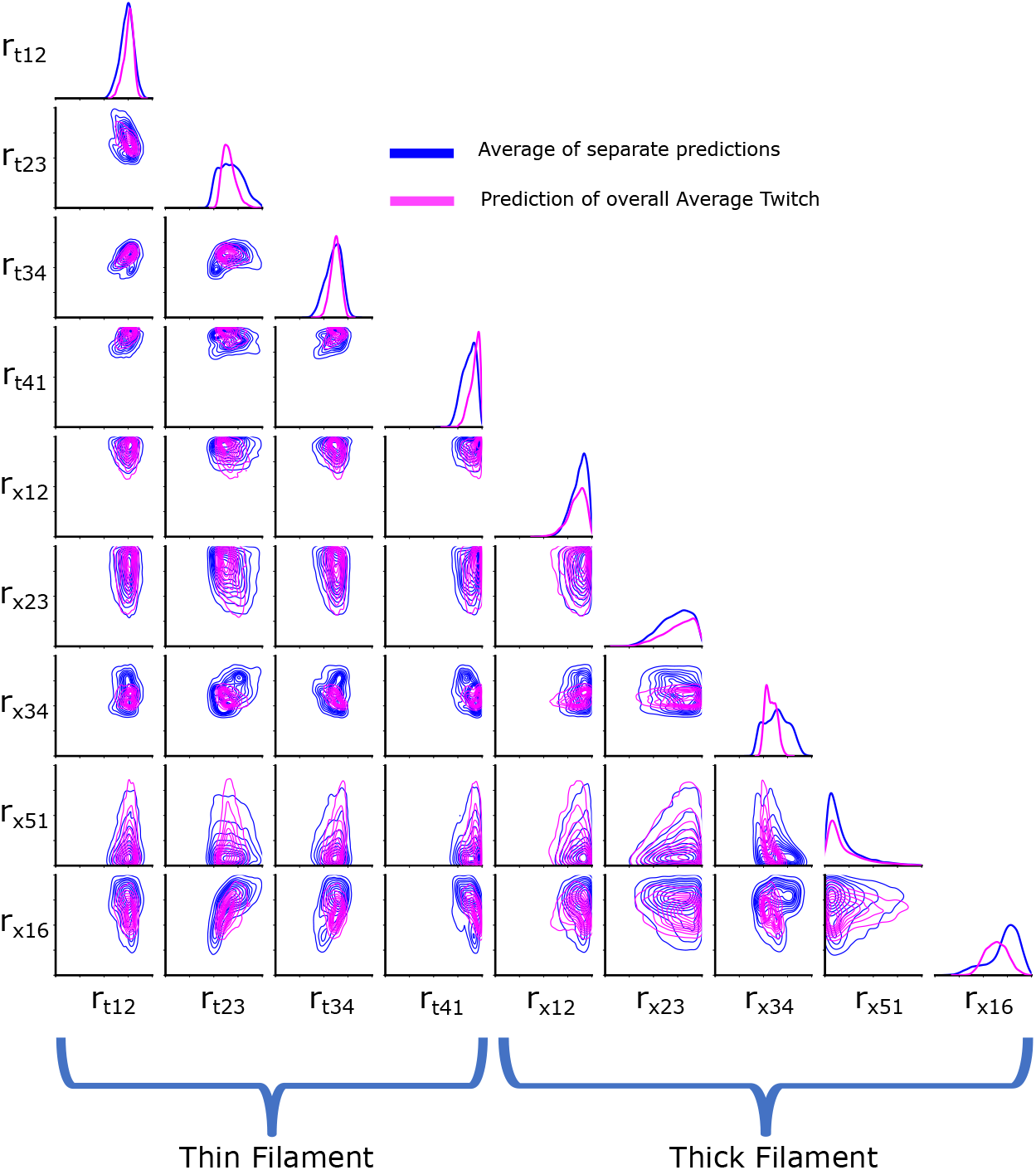
Blue indicates the distribution obtained by predicting the rate distribution of each of the individual experimental twitches separately, and then averaging the results. Pink indicates the prediction obtained by first averaging the separate twitches and predicting on that target alone. All predictions were done on the control twitch set.

### Time Step Convergence

**Supplementary Figure 3:**
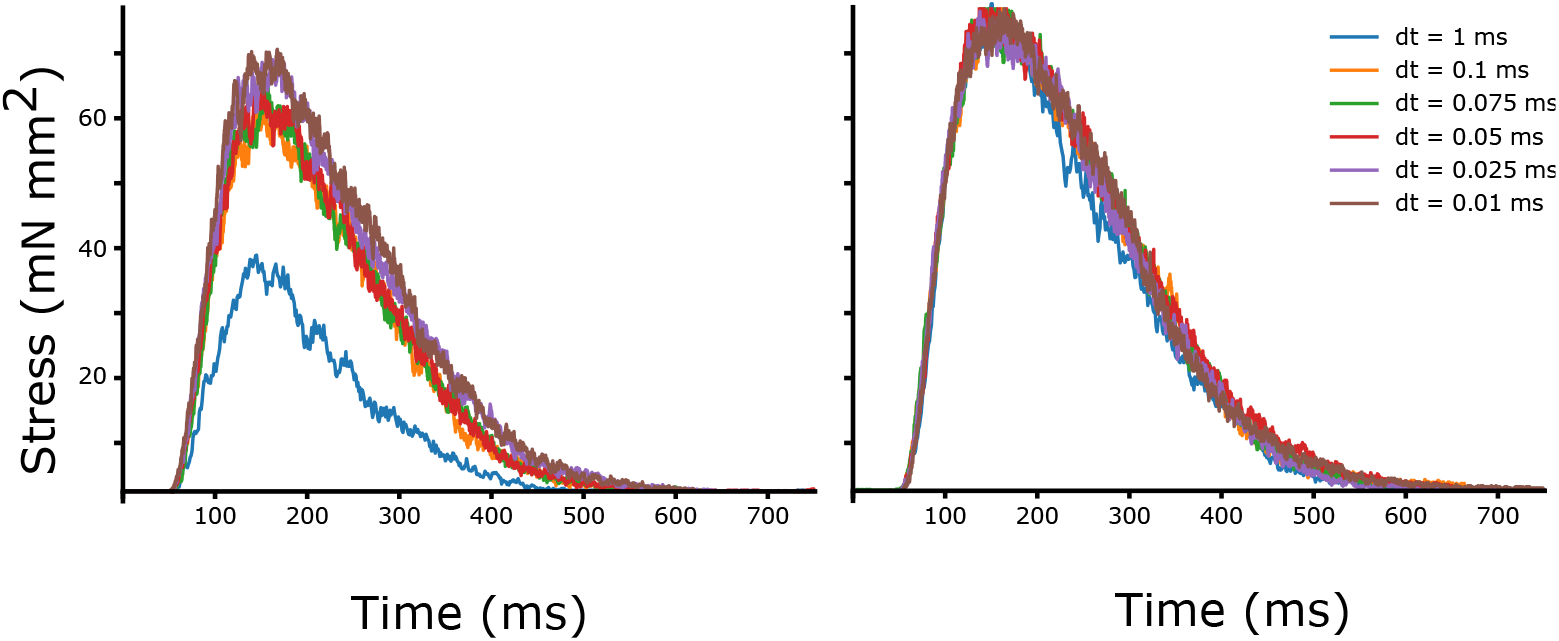
To show that the probability calculations converge faster, we simulated twitches at various time steps. Left uses the old numerical scheme, and shows that even very small time steps still have not converged. Right simulations use the numerical scheme in Eq. 1, and shows approximately the same twitches regardless of time-step used.

### Heterozygous vs Homozygous

**Supplementary Figure 4:**
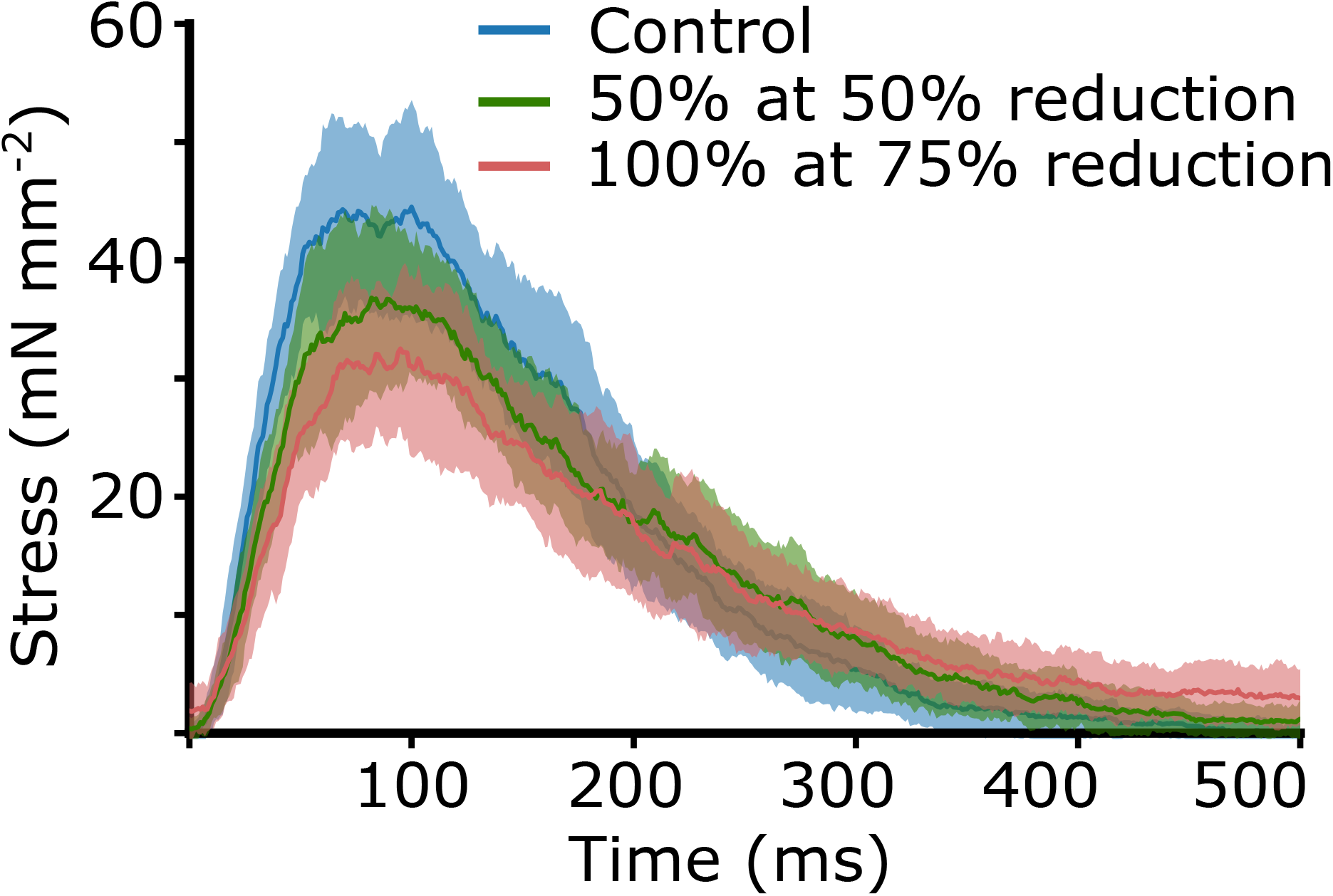
Because our muscle model is spatially explicit, meaning each crossbridge and binding site are treated individually, we can modify populations of crossbridges to simulate heterozygous genetic variants. Here we show a control simulation, using the same rate constants as the Control in Fig. 7. We then performed simulations in which *r*_*t*,12_ and *r*_*t*,14_ had been reduced by 50% for 50% of the binding sites (teal), and another simulation in which 100% of binding sites had *r*_*t*,12_ and *r*_*t*,14_ reduced by 25% (brown), therefore giving the same overall average reduction of 25%. But because of the spatially explicit nature of the model, the simulations result in different twitches.

### Kolmogorov-Smirnov test

**Supplementary Table 1.** The Kolmogorov-Smirnov test is used to show how similar or dissimilar two probability distributions are. The KS statistic shows the maximum difference (supremum) between two cumulative distribution functions (CDFs), and the KS Location indicates the rate factor at which the supremum occurs. The sign indicates which CDF was larger, with negative indicating that the value of the CDF for the control type was smaller, consistent with the bulk of the probability distribution’s mass occurring at larger rate factors, and positive indicating the opposite. The *p-*value in each pair-wise comparison was less than 10^−5^.

|  | Control and I61Q cTnC |  |  | Control and Danicamtiv |  |  |
| --- | --- | --- | --- | --- | --- | --- |
|  | KS Statistic | KS Location | Sign | KS Statistic | KS Location | Sign |
| $r_{t,12}$ | 0.51 | -0.25 | -1 | 0.12 | -0.79 | -1 |
| $r_{t,23}$ | 0.29 | -0.011 | 1 | 0.06 | -0.55 | -1 |
| $r_{t,34}$ | 0.43 | 0.60 | 1 | 0.07 | -0.08 | -1 |
| $r_{t,41}$ | 0.49 | 0.19 | -1 | 0.13 | 0.99 | -1 |
| $r_{x,12}$ | 0.37 | 0.76 | -1 | 0.02 | -0.78 | 1 |
| $r_{x,23}$ | 0.31 | 1.26 | -1 | 0.06 | -0.34 | -1 |
| $r_{x,34}$ | 0.36 | -0.24 | 1 | 0.31 | -0.65 | -1 |
| $r_{x,51}$ | 0.21 | -0.33 | 1 | 0.08 | 0.98 | 1 |
| $r_{x,16}$ | 0.84 | -0.75 | 1 | 0.48 | -0.15 | -1 |

### Validation Size

**Supplementary Figure 5:**
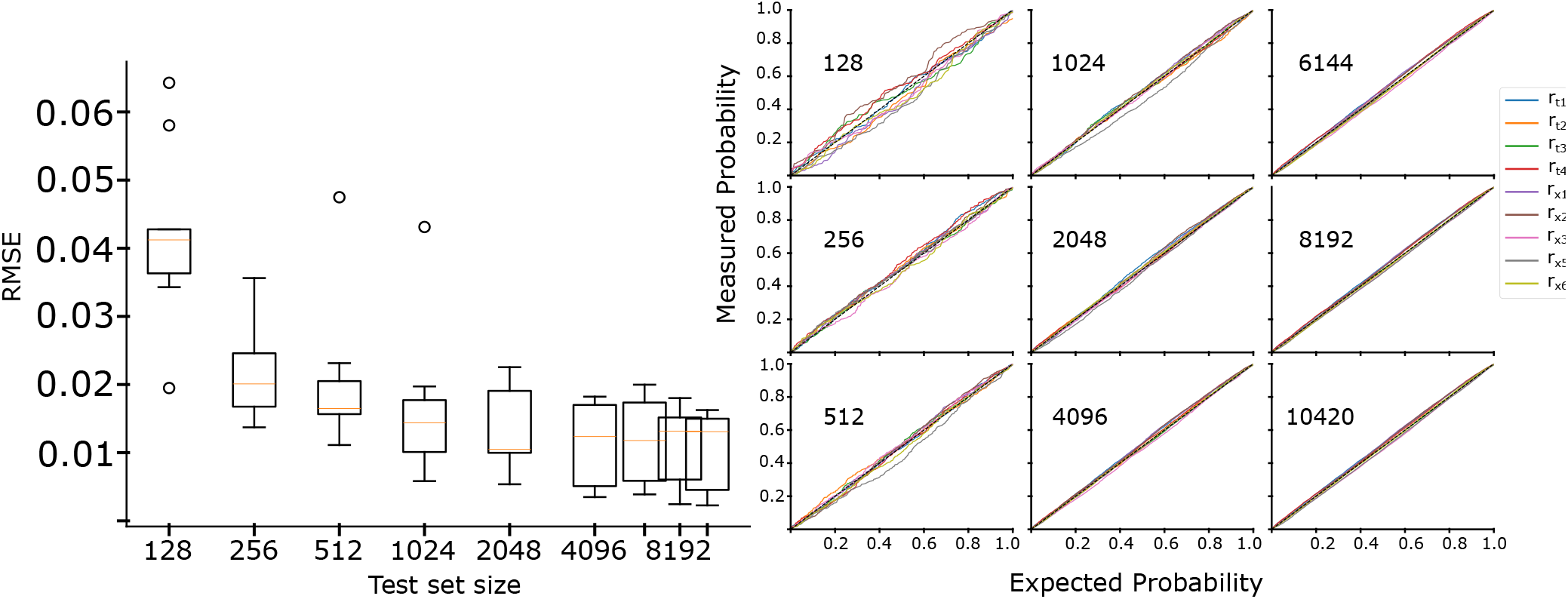
Here we plot the RMSE between the ideal and measured probability for a parameter to be in a certain region of the estimated probability distribution for test sets various sizes (Left). The box plots show the median (orange) and interquartile range. Note that the x-axis is plotted on a log_2_ scale. Right shows the probability-probability plots corresponding to each test set size for each of the rates we explored (indicated in the legend to the left of the graph). Also note that, above a test set size of 1000, there is convergence to a very low error in the probability estimate our method produces. As a reminder we used a test set of 11000, 1 % of the data set size

